# Tracking SARS-CoV-2 variants of concern in wastewater: an assessment of nine computational tools using simulated genomic data

**DOI:** 10.1101/2023.12.20.572426

**Authors:** Steven G. Sutcliffe, Susanne A. Kraemer, Isaac Ellmen, Jennifer J. Knapp, Alyssa K. Overton, Delaney Nash, Jozef I. Nissimov, Trevor C. Charles, David Dreifuss, Ivan Topolsky, Pelin I. Baykal, Lara Fuhrmann, Kim P. Jablonski, Niko Beerenwinkel, Joshua I. Levy, Abayomi S. Olabode, Devan G. Becker, Gopi Gugan, Erin Britnell, Art F.Y. Poon, Renan Valieris, Rodrigo D. Drummond, Alexandre Defelicibus, Emmanuel Dias-Neto, Rafael A. Rosales, Israel Tojal da Silva, Aspasia Orfanou, Fotis Psomopoulos, Nikolaos Pechlivanis, Lenore Pipes, Zihao Chen, Jasmijn A. Baaijens, Michael Baym, B. Jesse Shapiro

## Abstract

Wastewater-based surveillance (WBS) is an important epidemiological and public health tool for tracking pathogens across the scale of a building, neighbourhood, city, or region. WBS gained widespread adoption globally during the SARS-CoV-2 pandemic for estimating community infection levels by qPCR. Sequencing pathogen genes or genomes from wastewater adds information about pathogen genetic diversity which can be used to identify viral lineages (including variants of concern) that are circulating in a local population. Capturing the genetic diversity by WBS sequencing is not trivial, as wastewater samples often contain a diverse mixture of viral lineages with real mutations and sequencing errors, which must be deconvoluted computationally from short sequencing reads. In this study we assess nine different computational tools that have recently been developed to address this challenge. We simulated 100 wastewater sequence samples consisting of SARS-CoV-2 BA.1, BA.2, and Delta lineages, in various mixtures, as well as a Delta-Omicron recombinant and a synthetic “novel” lineage. Most tools performed well in identifying the true lineages present and estimating their relative abundances, and were generally robust to variation in sequencing depth and read length. While many tools identified lineages present down to 1% frequency, results were more reliable above a 5% threshold. The presence of an unknown synthetic lineage, which represents an unclassified SARS-CoV-2 lineage, increases the error in relative abundance estimates of other lineages, but the magnitude of this effect was small for most tools. The tools also varied in how they labelled novel synthetic lineages and recombinants. While our simulated dataset represents just one of many possible use cases for these methods, we hope it helps users understand potential sources of noise or bias in wastewater sequencing data and to appreciate the commonalities and differences across methods.

## Introduction

Wastewater-based surveillance (WBS) is a powerful tool to track pathogens, genes, and drugs across the scale of a building, city, or region^1^. The ongoing COVID-19 pandemic has shifted the attention to pathogen surveillance^2^, and over the course of the pandemic, it has become increasingly common to track SARS-CoV-2 in wastewater^3^. SARS-CoV-2 can be tracked in wastewater due to infected individuals shedding viral particles of SARS-CoV-2 in faeces^4–7^. WBS has become a vital component of SARS-CoV-2 surveillance and is implemented across much of the world^3^.

WBS of SARS-CoV-2 often relies on qPCR-based quantification of viral RNA – providing a rapid estimate of infection rates in the population^8^. A limitation of qPCR is that it provides little to no information about viral diversity – including the presence and relative abundance of genetically distinct viral lineages, including variants of concern (VOCs). Designing primers to distinguish among VOCs by qPCR^9^ can be laborious and is hindered by novel viral mutations^10^. A complementary approach to qPCR is to measure viral diversity by sequencing parts, or the entirety, of the viral genomes found in wastewater^11–15^.

Sequencing wastewater provides an opportunity for cost-effective epidemiological surveillance^16^. However, challenges persist both upstream and downstream of sequencing. Upstream, the quality of results may vary according to the population size contributing to the wastewater catchment area^8^, variability in viral shedding^17,18^, virus enrichment methods^19^, and in-sewer degradation of viral-particles (e.g. pH, temperature, travel-time)^20,21^. These and other factors influence how much viral RNA is collected for sequencing^22^. Typically, viral RNA recovered from wastewater is at a lower concentration and more degraded than in clinical samples, especially outside major waves of infection^23^. The combination of low RNA concentrations and PCR inhibitors in wastewater can make amplicon-based sequencing methods prone to uneven coverage across the genome (*e.g.* due to some amplicons failing to amplify) or outright failure. Downstream of sequencing, mixtures of diverse viral lineages from wastewater can make it difficult to correctly assign mutations on short reads to the correct lineage^24,25^. High viral diversity in wastewater can lead to consensus genomes which are prone to chimeras and errors, and mostly represent the dominant lineage while failing to identify rarer ones^21^.

A variety of computational tools have been developed recently to address the challenges of wastewater sequencing, with the common goal of inferring the presence and (in most cases) the relative abundances of various SARS-CoV-2 lineages present in wastewater. In this benchmark, we focus on SARS-CoV-2 whole-genome sequencing using short reads from tiled amplicons to identify specific viral lineages and their relative abundances^26^. This approach is currently more commonly used than long-read sequencing^27^. In this study, we compared the performance and output of nine commonly used computational tools, available between winter 2021 and spring 2022 in a semi-blind benchmark on simulated wastewater sequencing data. These tools differ in the precise statistical methods, criteria for inferring lineages, as well as the databases used (Table 1). The benchmark dataset consisted of simulated *in silico* mixtures of SARS-CoV-2 genomes belonging to BA.1, BA.2, Delta and a Delta-Omicron recombinant, as well as a synthetic “novel” lineage (Methods).

**Table 1.** Summary of nine approaches included in this study. Input required for each approach is either FASTQ (sequencing reads), or VCF (allele frequency table relative to the Wuhan reference). The database used by each approach was divided into two categories: Curated Mutation Profiles, where a selection of alleles present per lineage is done by a functional or phylogenetic criteria, or Selected Alleles from GISAID Genomes, where all or some GISAID genomes are aligned to the reference genome, and alleles are selected based on allele-frequency. For VLQ this is used in selecting a reference set for lineages when performing pseudoalignment. Database availability refers to how users can access the database, by accessing the tools’ Github repository or by executing a command within the tool.

| Tool | Package Manager | Dependencies | Input | Method to estimate lineage relative abundances | Database Type | Database Details | Database Availability | Reference |
| --- | --- | --- | --- | --- | --- | --- | --- | --- |
| Alcov | Pip | Cutadapt, minimap2 Python Packages: Fire, numpy, pandas, scikit-learn, matplotlib, seaborn, pysam | FASTQ | Linear Model: Ordinary Least Squared | Curated Mutation Profiles | Cov-Spectrum | Github | 28 |
| Basic | NA | blast+, PRINSEQ, fastp, bwa, picard, samtools, varscan, python3, R | FASTQ | Linear Model: Constrained Linear Model | Selected Alleles from GISAID Genomes | Allele-frequency >= 90% | Github | 30 |
| Freyja | Conda | iVar, samtools, USHER Python Packages: cvxpy, numpy, pandas | FASTQ | Constrained (weighted) Least Absolute Deviations | Curated Mutation Profiles | Marker mutations from USHER global phylogenetic tree | Github or; Via Freyja | 10 |
| Gromstole | NA | cutadapt, minimap2, Python, R | FASTQ | Quasibinomial Regression Model | Curated Mutation Profiles | Cov-Spectrum | Github | 31 |
| LCS | NA | Snakemake, samtools, GATK4, bwa, picard, USHER Python Packages: biopython, pysam, pyvcf, pandas, cvxpy | FASTQ or VCF | Log-likelihood Maximization Model | Selected Alleles from GISAID Genomes | Allele-frequency >= 80% with phylogenetic verification | Github or; Via LCS | 32 |
| Lineagespot | bioconductor | R | VCF | Average Allele Frequency | Curated Mutation Profiles | Outbreak.Info or Pangolin | NA | 33 |
| VLQ | NA | kallisto, samtools, minimap2, bwa, bbmap Python Packages: pyvcf, pysam | FASTQ | Pseudoalignment to reference genomes (Kallisto) | Selected Alleles from GISAID Genomes | Reference genomes selected to capture alleles with frequency >= 50% | Via VLQ | 35 |
| V-pipe | Conda | COJAC, ShoRAH, LolliPop Python Packages: pysam, pandas, numpy, pyyaml, strictyaml, requests, click, poetry-core | FASTQ | Linear Model: Constrained Linear Model (LolliPop) and Dirichlet Process Mixture Model (ShoRAH) | Curated Mutation Profiles | Cov-Spectrum and UKHSA Genomics Public Health Analysis Variant-Definitions | Via COJAC | 37,38,39,40 |
| Pipes | NA | bowtie2, R (ape) | FASTQ | Expectation-Maximization Algorithm | Selected Alleles from GISAID Genomes | Calculated phylogenetic internal nodes for each lineage | Via Pipes | 34 |

The goal of the simulated dataset was to challenge the tools with respect to their lineage identification and quantification under variable conditions (i.e., amplicon drop-outs, varying depth of coverage and read length, and in the presence of “background noise”). These conditions are meant to reflect real-world financial and operational constraints on sequencing methods and effort. While the dataset is meant to reflect these considerations, it remains a model use case that does not necessarily reflect the complexity of real-world data. We also acknowledge that this is not a comprehensive comparison of all available approaches, nor can this be considered a fully quantitative or unbiased comparison, not least because the developers of each tool were given the benchmark dataset and ran their tool as they saw fit, according to their own best practices. We found that in general all tools performed their function very well and that their results were largely robust to such challenges as low sequencing depth or shorter read lengths, thus raising confidence in VOC calling, no matter the specific sequencing conditions and tool used. The tools included in the benchmark test are described in detail in their respective publications. A brief description of each tool follows.

### Alcov

A python package designed to leverage both unique, lineage-specific mutations, and shared, non-lineage specific mutations^28^. “Double-counting” of shared-mutations is avoided by performing a lineage prediction *a priori* to estimate expected mutation frequencies in each sample. With an expected, and observed mutation frequency, a most likely frequency is determined by minimising the error between predicted and expected mutation frequencies. For lineages in which there are at least 100 sequences, Alcov uses lineage-defining mutations selected from cov-spectrum^29^ that are present in >90% of lineages.

### Basic

A pipeline devised to analyse wastewater sequences from variant lineages during WBS in Québec between March 2020 and July 2021^30^. The pipeline is composed of two components: variant calling of single nucleotide variants that pass coverage and quality thresholds, followed by post-variant-calling analysis. Post-variant-calling calculates relative abundances of lineages with a constrained linear model to fit the lineages’ signature mutations (at least three per lineage) at allele-frequencies over 90%. This pipeline was designed and tested before the first Omicron wave. It also only considers minor alleles in wastewater samples at a frequency of 25% or higher, making it less sensitive to rare variants and more attuned to confidently identifying common lineages.

### Freyja

A suite of analysis tools for real-time wastewater genomic surveillance. Freyja includes a “demix” function that enables users to estimate the relative abundance of virus lineages in a mixed sample, such as wastewater^10^. This method encodes genomic sequencing of virus mixtures using single nucleotide variant (SNV) frequencies and leverages unique mutational barcodes for each lineage (assembled using the most recent UShER global phylogenetic tree). To recover relative lineage abundances from a mixture, the barcodes and SNV encodings are integrated in a mixture model formulation that is solved using a constrained, depth-weighted least absolute deviations approach. The use of depth weighting confers robustness to limited sequencing coverage, a common feature of wastewater samples. Freyja is written in Python, has a command line interface, and is available via conda.

### Gromstole

A lightweight pipeline^31^ that consists of Python and R scripts to preprocess samples and extract the read coverage and mutation frequencies per nucleotide site for each sample. It uses a curated list of lineage-specific mutations. The frequency of a lineage in the sample is estimated by assuming the frequency of each associated mutation is an independent outcome. A quasibinomial regression model is used to compute 95% confidence intervals for each frequency estimate. The pipeline is designed to be lightweight and minimise the number of dependencies required. It was not designed to call novel variants or recombinant forms of SARS-CoV-2. To mitigate the problem of numerous shared mutations among lineages, the tool uses a curated list of lineage-specific mutations to exclude mutations that occur at substantial frequencies in other (“off-target”) lineages, with the concession of substantially reducing the number of mutations that can be used to estimate lineage frequencies.

### LCS

A viral genome deconvolution approach based on relative frequencies of polymorphisms found in known SARS-CoV-2 variants^32^. The reference database is built, first, with pre-defined SARS-CoV-2 lineages assigned to variant groups (VGs) according to the currently tracked variants defined by the World Health Organization. Next, manually curated genome designations from the Pango Lineage Designation Committee are assigned to the corresponding VGs. These genomes, whose sequences are available in the GISAID database, are mapped to the SARS-CoV-2 reference genome and polymorphic sites with allele frequency greater than 80% in at least one VG are selected. Based on these markers, the method fits a mixture model to pools of SARS-CoV-2 samples, considering the relative frequencies of polymorphisms found in each pool to obtain a maximum *a posteriori* estimate of the relative contributions of SARS-CoV-2 variants to the pool.

### Lineagespot

A bioconductor R-package^33^ that uses VCF files, which contain all the nucleotide (and corresponding amino acid) changes identified in a sample, along with a file containing all lineage-assignment mutations. Lineagespot uses identified mutations grouped and assigned to the respective SARS-CoV-2 lineages and computes a number of metrics (such as the average allele frequency, and the minimum non-zero allele frequency). It attempts to determine the presence of a specific lineage (specified as input to the tool) by calculating the likelihood of its existence in the sample, rather than its relative abundance. This benchmark exercise, which assesses the ability of each tool to infer lineage relative abundances, therefore extends beyond the original scope of Lineagespot.

### Pipes *et al* 2022 (“Pipes”)

A method^34^ that builds a reference database, constructs a read × haplotype mismatch matrix, and then uses an Expectation-Maximization (EM) algorithm to obtain the maximum likelihood estimates of the proportions of different haplotypes in a sample. The reference database is constructed using a phylogenetic imputation approach.

### Viral Lineage Quantification (VLQ)

A pipeline^35^ that differentiates between lineages and sub-lineages based on a reference set that can be tailored to the location and time of sampling. VLQ consists of three steps: first, it constructs a reference set consisting of one or more full genome sequences per SARS-CoV-2 lineage of interest, selected to represent local variation. Then, VLQ uses kallisto^36^, an algorithm for RNA transcript abundance quantification from RNA-Seq data, to predict the relative abundance of each reference sequence. Finally, VLQ performs basic post processing: background noise is removed by filtering out predictions below a user-specified minimal abundance level and subsequently the predicted abundance for sequences from the same lineage (or VOC) are summed to obtain the total abundance per lineage (or VOC).

### V-pipe

In the context of this benchmark, V-Pipe refers to a workflow developed in the Beerenwinkel Lab at ETH Zurich^37^ which includes components developed for wastewater sequencing data analysis: COJAC^38^, LolliPop^39^ and ShoRAH^40^. V-pipe preprocesses samples, and COJAC searches for co-occurrences of variant signature mutations on individual PCR amplicons. For the detected variants, relative abundances are quantified using LolliPop by simultaneously smoothing and deconvolving the observed mutation frequencies in sequencing reads into relative abundances of the variants and providing confidence bands. ShoRAH identifies unknown lineages by a Bayesian nonparametric read clustering method which denoises the sequencing data and reconstructs local haplotype sequences. Inferred local haplotypes with mutations different from any known variants are considered signatures of new variants.

## Results

The benchmark test dataset consisted of 100 simulated wastewater samples containing either single lineages or mixtures, at high or low coverage, and with long or short read lengths (Supplementary Table 1). Samples were provided to the developers of each tool and run according to their own best practices on their own systems. Each team was aware that samples would contain BA.1, BA.2 and Delta sequences, but were not told the composition of each sample. We evaluated how each tool identified lineages and how well they estimated relative abundance.

We first considered samples that contained only a single lineage of either BA.1, BA.2, or Delta, at low coverage (37.5x mean depth) and short 150 bp reads (**Figure 1**). Most of the tools correctly identified the true lineage, at close to the expected 100% frequency (**Figure 1**), with the exception of Lineagespot, which is designed to identify the presence/absence of a variant rather than the abundance (where it provides only the absolute minimum guaranteed level of presence). False positive lineages were sometimes inferred, typically at low frequency. The exception was the Basic approach which often could not distinguish BA.1 and BA.2 (**Figure 1**).

**Figure 1.**
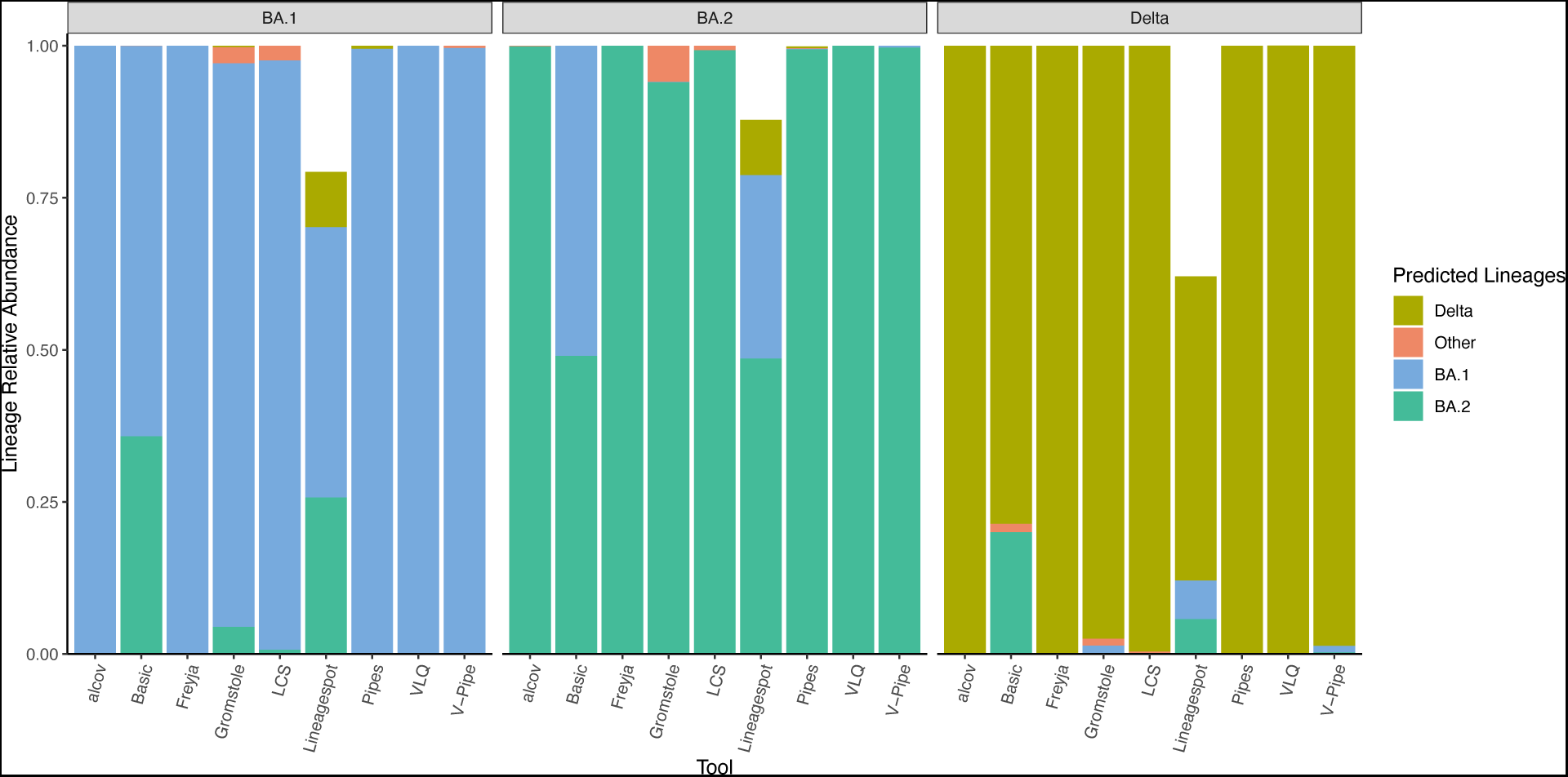
Inference of BA.1, BA.2 or Delta in single-lineage samples. Relative abundance of single-lineage predictions for samples of BA.1, BA.2, Delta with simulated 150 bp reads and low coverage (37.5x mean depth). ‘Other’ is defined as lineage calls that are not either BA.1, BA.2 or Delta.

Next, we considered mixed-lineage samples and compared the ability of each tool to identify the correct lineages (F1 score) and estimate their relative abundances (Root-mean-square error; RMSE). With some variation, tools were generally able to identify the correct lineages present (**Figure 2A**) and estimate their relative abundances (**Figure 2B**). Most tools had consistently high F1 scores and low RMSE scores. Exceptions were lineagespot and Basic, which had lower F1 scores and higher mean RMSE across simulation parameters. Lineagespot’s higher RMSE is likely because relative abundance estimation is beyond the intended scope of the tool, and estimated relative abundances often totalled to below 100%. Basic’s result can be explained by the fact that the approach was designed and tested on lineages pre-Omicron, and for detection of abundances above 5%.

**Figure 2.**
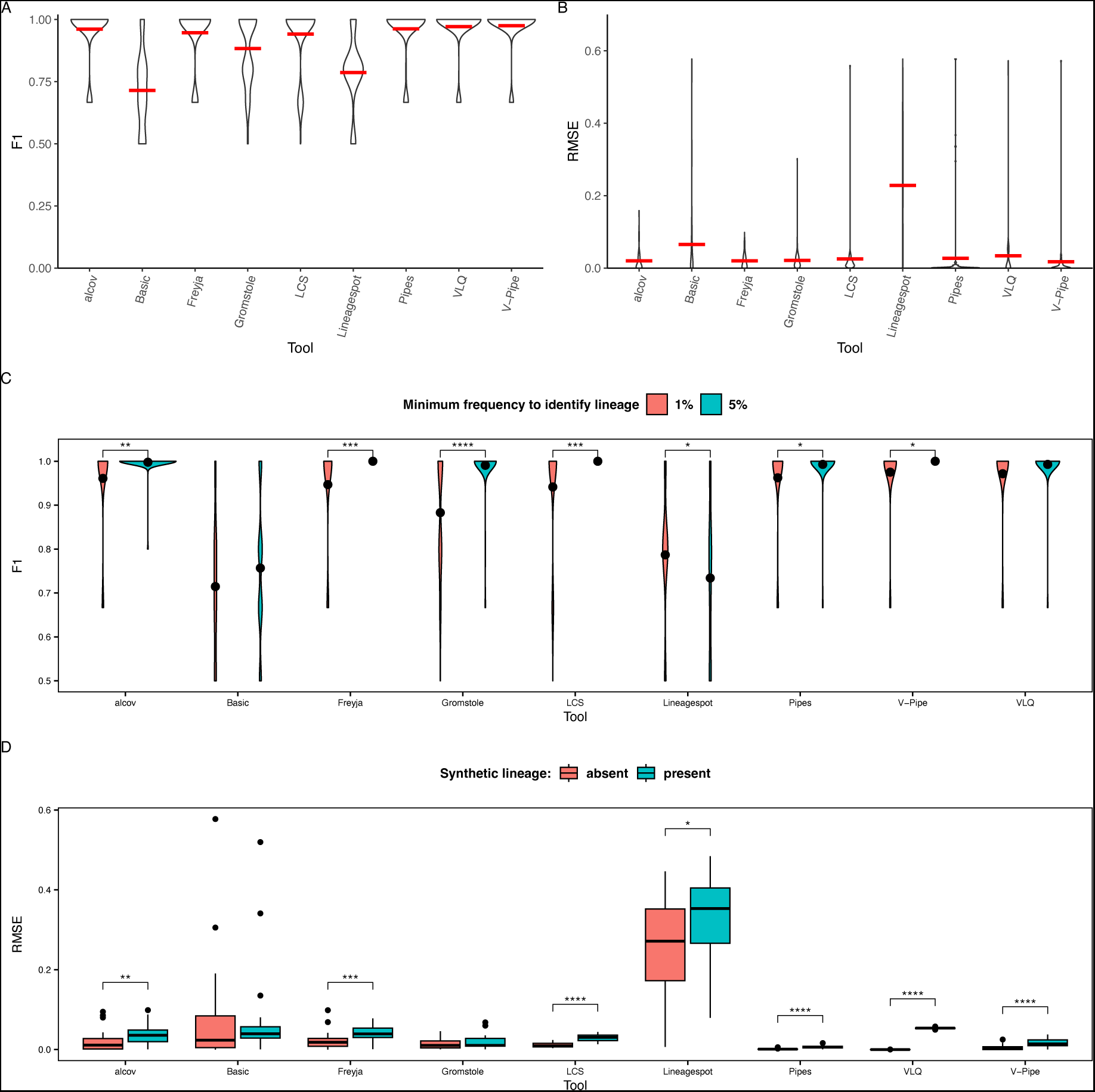
Lineage identification and quantification in single- and mixed-lineage samples. Based on all 100 simulated samples, we calculated (A) F1 scores and (B) Root-mean-square error (RMSE) for each tool. Horizontal red lines show the mean. Lineage presence is defined by detection at ≥ 1% abundance in a sample. (C) Comparison of F1-scores with lineage presence defined by ≥ 1% or ≥ 5% abundance (black point shows the mean F1). (D) Comparison of relative abundance estimates, RMSE, between mixed-lineage samples with a synthetic lineage present (n=22) or absent (n=36). * wilcox p-adjusted (BH) < 0.05

Lineage detection in a sample is intuitively more likely if the lineage is present at a higher frequency. In our example, a lineage was considered present in a sample if it was found at 1% or more. However, a previous study set the limit of detection at 5%^10^. We compared F1 scores at 1% vs. 5% cutoff for detection, and found that most tools had higher F1 scores when the 5% cutoff was used (**Figure 2C**). We also compared lineage identification and relative abundance quantification between samples with different parameters: high or low coverage, 150 or 250 bp reads, with or without amplicon drop-outs, and with or without background noise (i.e., in the presence of synthetic lineage). Sequencing always comes with a background error rate, and amplicon-based sequencing can be sensitive to amplicon dropouts when mutations interfere with primer binding. We found no significant differences between samples with and without amplicon dropout, low-coverage, or short-reads in lineage identification or estimates of relative abundances. However, the presence of background noise had a significant effect on lineage quantification (**Figure 2D**). Adding background noise, in the form of a synthetic, unknown lineage, increased the RMSE consistently across tools. The magnitude of this effect was relatively small, but could impact lineage deconvolution in more complex background conditions. Together, these findings highlight that lineage calling can be confidently performed even with relatively low sequencing depth and short read lengths, but that lineages predicted to be present below 5% relative abundance should be interpreted with caution.

Having explored the performance of tools within their intended use of detecting known lineages, we considered more challenging tasks of identifying unknown or recombinant lineages. First, we asked how the tools interpret a synthetic lineage which is not present in any of the databases. We simulated this synthetic lineage based on the Wuhan reference genome background, adding 50 mutations previously observed in real genomes, but never rising to high frequency nor appearing together in the synthetic combination (Methods). This lineage could represent a newly evolved or introduced lineage. Most tools (Alcov, Basic, Gromstole, LCS, Lineagespot, V-Pipe) identified the reads as belonging to ‘Other’ lineages rather than attempting a designation such as BA.1, BA.2, or Delta (**Figure 3A**). Freyja, which employs the UShER phylogenetic tree for lineage-identification, placed the synthetic lineage near the root as a deep-branching ‘B’ lineage (**Figure 3A**). There were also some instances of false positive identification of Delta or BA.1 (Pipes and VLQ) (**Figure 3A**). Together with the observation that the synthetic lineage can weakly but consistently add error to relative abundance estimates of known lineages (**Figure 2D**), these results highlight the challenges associated with handling novel lineages in mixed wastewater samples.

**Figure 3.**
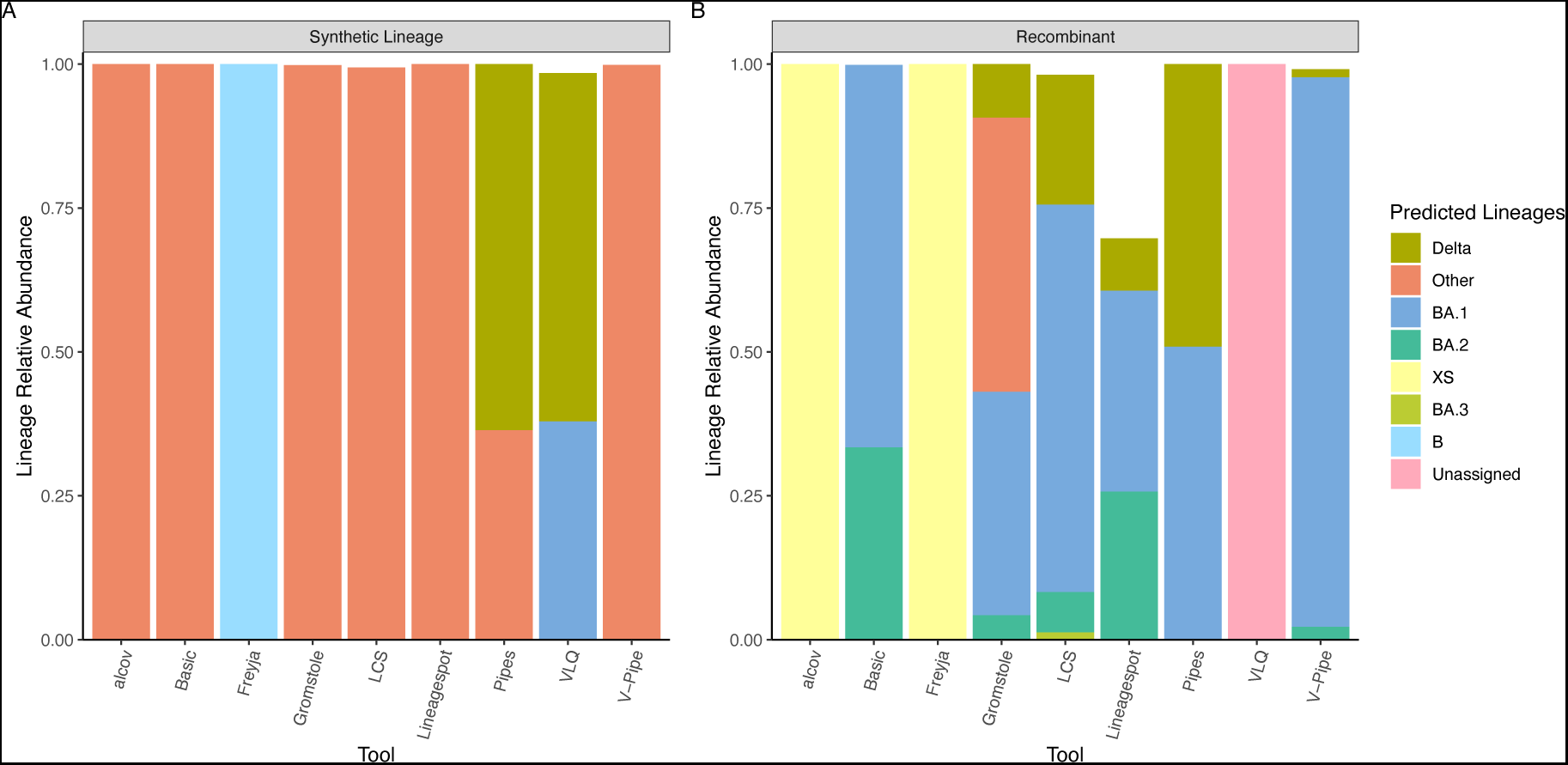
Identification of unknown lineages in simulated wastewater sequences. (A) Each tool was applied to a sample containing a simulated synthetic lineage sequenced with 150 bp reads and low coverage (<50x). The synthetic lineage is the Wuhan reference genome with the addition of 50 non-lineage defining mutations. (B) Each tool was applied to a sample containing a simulated recombinant of Delta:BA.1.1 from low-coverage 250 bp reads. Only predictions of lineages >1% are represented in the barplot.

Second, we explored how each tool dealt with a recombinant of Delta and BA.1.1. Several tools (Pipes, Gromstole, LCS, Lineagespot, Basic, and to a lesser extent V-Pipe) split their lineage calls between the parent lineages, Delta and Omicron (**Figure 3B**). Freyja and Alcov called it ‘XS’, which is a Delta:BA.1 recombinant, while VLQ left it unassigned.

## Discussion

Prior to the SARS-CoV-2 pandemic, wastewater-based surveillance had been an important part of the epidemiological polio surveillance and eradication effort^41^. The discovery that one could differentiate between vaccine and non-vaccine derived polio virus without culturing the virus from environmental samples^42^ showed that it should also, in principle, be possible to detect and estimate the relative abundances of different lineages of SARS-CoV-2 from wastewater^43^. Detecting and quantifying viral lineage relative abundance from a mixed wastewater sample is greatly facilitated by clinical genomic protocols^44,45^; however, the processing of this sequence data is not trivial, providing the motivation for this study.

Nine different teams, across six different countries, developed distinct approaches to distinguish SARS-CoV-2 lineages from wastewater. We acknowledge that this was not an exhaustive set of methods, and new methods have been developed since our project was initiated. The common goal of all approaches is to identify SARS-CoV-2 lineages and estimate their relative abundance. Lineagespot, the exception, was not designed to infer the relative abundances directly. Each team made assumptions during the development of their approach as to how, and when, their tool would be used, and this benchmark did not aim to reflect all possible scenarios. For example; should all SARS-CoV-2 genomes circulating globally be used in the set of lineage-defining alleles, or only a locally/temporally-matched subset? Are lineage-defining alleles curated based on functional characteristics, phylogenetics, or frequency within the population? What type of sequence preprocessing is required, and/or optional? Should all alleles observed in a lineage be included? If not, what criteria should be used in selecting lineage-defining alleles? What priorities should be favoured in selecting a model of lineage deconvolution for mixed-viral samples? Our simulated benchmark dataset represents one common public health use case^46^, and addresses technical considerations upstream of bioinformatic analysis (i.e., read length, and minimum genome coverage), but falls far short of evaluating all use cases for the individual tools.

This study was initiated when most of the approaches were still under active development, which made running each tool locally by one user impractical. Instead, the benchmark was implemented as a semi-blind trial: each team knew to expect a mixture of Delta and Omicron lineages, but not a synthetic ‘novel’ sequence or a simulated recombinant genome. Each team ran their own method on the simulated data and was thus able to make some ‘expert’ decisions that might not be apparent to a naive user. For this reason, we discourage side-by-side comparisons between the tools. A more stringent comparison would have one group run all the methods independently, thus more closely simulating the experience of a new user running a tool ‘out of the box.’ Such an approach would allow a more direct measure of user-friendliness, reproducibility, and compute time.

Overall performance was tested based on the ability to identify lineages and estimate their relative abundance. Lineage identification did not involve distinguishing finer-grained PANGO-lineages (e.g., BA.2.1.7 vs BA.2.23.1), a feature not supported by all approaches. For the sake of comparison, we focused on broader-scale distinctions among WHO-designated VOCs (e.g., Delta vs Omicron) or Nextstrain Clades (e.g., BA.1/21K vs BA.2/21L). To allow for comparisons between approaches, we collapsed finer-scale lineage predictions, if present, into broad categories of Delta, BA.1, and BA.2.

All approaches require the identification of lineage-defining alleles for lineage detection, and the steps involved vary among approaches – again making side-by-side comparisons challenging. We classified tools into two categories of reference databases: mutation lists curated by the team or based on the allele frequencies within a lineage in the GISAID database. Genomes in the benchmark came from multiple different countries, which means that approaches that build reference sets based on a subset of genomes from GISAID (e.g., VLQ) could not leverage the metadata (i.e., time or geographic location) for selecting suitable reference sequences per lineage but instead included the reference genomes used in building the benchmark dataset and others present in GISAID.

We found that most approaches identify lineages and estimate their relative abundances as intended. A surprising finding was that tools could detect lineages at abundances as low as 1%. However, increasing the cut-off for detection to 5% reduced false-positives, and significantly improved F1 scores for most tools, so detection below 5% should be interpreted with caution. We also explored several factors that could impact the implementation of WBS: genome coverage, read-length, and amplicon drop-out. None of these factors significantly affected lineage calling or relative abundance estimates. Therefore, reducing coverage and read length (to a point) could help cut costs while preserving most of the desired lineage information. These considerations could help make WBS epidemiologically informative and cost-effective, particularly in resource-poor settings^47^.

With the goal of testing how tools dealt with a novel lineage, we simulated a synthetic lineage with non-lineage defining mutations. As none of these mutations are lineage-defining, this could also be viewed as simulating recent *de novo* mutations in the population, or common sequencing errors. Most tools (Alcov, Basic, Freyja, Gromstole, LCS, Lineagespot, Pipes, V-Pipe) could correctly identify the presence of a non-Omicron, non-Delta lineage in the sample. The presence of a synthetic lineage, or background sequencing noise in mixed samples, weakly but significantly impacts the relative abundance estimation for a number of tools (Alcov, Freyja, LCS, Lineagespot, Pipes, VLQ, and V-Pipe). We also investigated how tools handled an unclassified recombinant of Delta-Omicron. Faced with an unclassified recombinant genome, tools either identified the two parental lineages, acknowledged it was an unknown lineage, or found the closest known lineage. While recombinant and synthetic lineage identifications fall beyond the intended scope of most tools, they give us some idea how tools might behave when faced with the diversity of lineages present in a real wastewater sample.

The simulated dataset described here was developed in 2021 and the nine teams applied their methods to the data in 2022. This study therefore reflects the questions we had at the time. Since then, the recombinant XBB (Nextstrain clade 22F) has risen to global abundance, and convergent mutations have made it increasingly challenging to disentangle lineages. As we write, SARS-CoV-2 genome sequencing has been deployed in wastewater from across the globe (e.g., USA, Canada, Uruguay, Japan, Italy, India, Spain) to augment surveillance efforts during Gamma, Delta, and Omicron (BA.1 to XBB/BQ) waves^12,13,25,26,48^. The uptake of wastewater sequencing data to bring lineage-level information to WBS highlights the importance of benchmarking and developing standards. Such standards have emerged for clinical sequencing of SARS-CoV-2^49^ and we hope that our work will inspire further efforts towards standards for wastewater sequence data. Just as efforts to quantify and sequence SARS-CoV-2 from wastewater built on the knowledge gained through polio surveillance efforts, WBS is now being applied to other pathogens such as human influenza, metapneumovirus, parainfluenza, respiratory syncytial virus (RSV), and rhinovirus^50,51^. Wastewater has mostly been sampled and analysed in cities, but there are efforts to expand surveillance to airplanes^52^ and more rural communities^53^. As new sampling schemes, sequencing technologies, and computational methods are developed, it is essential to understand their behaviour using simulated data.

## Methods

### Benchmark Dataset

We randomly selected 10 high-quality genomes each belonging to Delta, BA.1 and BA.2 lineages from among all the genomes listed in the cov-lineages pango-designation lineages (github.com/cov-lineages/pango-designation/blob/master/lineages.csv?; accessed March 2022) and downloaded them from GISAID^54^ (Supplementary Table 2). The genomes were selected randomly, with no bias toward genomes that were ‘representative’ of their lineage. All genomes were verified not to contain long stretches of Ns. The recombinant was represented by four AY.119.2:BA.1.1 recombinants genomes^55^ (Supplementary Table 2). To create a synthetic genome, we started from the reference genome hCoV19/Wuhan/WIV04/2019, to which we added 50 mutations. These mutations were randomly selected from those present at minor frequencies (10% or less) in Alpha lineage genomes in the GISAID multiple sequence alignment published on 27-03-2022. To be chosen, mutations needed to be observed in at least 1000 genomes and were double-checked to ensure that they were not defining for any known PANGO constellation (https://cov-lineages.org/constellations.html). Mutations were excluded if they caused a stop codon or were located in the intergenic area of the genome.

Amplicons were created by cutting out the ARTIC v4.1-based fragments from the genome using seqkit^56^. To simulate uneven coverage across genomes we also created a BA.1 fragment set in which three PCR amplicons (6, 55, 74) were dropped from all BA.1 genomes. Artificial reads were created using insilicoseq^57^. We created custom error models based on wastewater samples sequenced with a read length of either 150 or 250 bp using bwa mem (v. 0.7.17) and samtools (v.1.12). The *in silico* reads were subsequently randomly subsetted and combined to create 100 different samples representing single strains, as well as mixtures of two or three strains with differing coverage and read lengths. Low coverage samples with 250 bp reads included 4000 reads; 150 bp read samples included 7500 reads. High coverage samples with 250 bp reads included 80,700 reads; 150 bp read samples included 150,000 reads. A total of 100 samples were generated (Supplementary Table 1).

### Tools

#### Alcov

Preprocessing for Alcov was done with the alcov-prep.py script which uses cutadapt (4.1), minimap2 (2.24.r1122), and samtools (1.7) to trim adapter sequences and map reads to the reference genome. Alcov was then run on each sample to generate lineage predictions using the command “alcov find_lineages --min_depth=10 --unique=False --csv=True samples.txt” with constellation files updated on June 20, 2023.

#### Basic

The variant calling step was run with default parameters: minimum coverage of 50x and a conservative minimum allele frequency of 25% based on findings previously reported^30^. Post-variant-calling was modified for this benchmark to include BA.1, BA.2, and BA.3, not included in the original pipeline, and updated the mutation prevalence file built using multiple-sequence-alignment on all genomes. Mutations with prevalence of >=90% for each VOC investigated here (Alpha, B.1.351, Gamma, Delta, Lambda, Mu, BA.1, BA.2, BA.3) based on the GISAID msa from 27-03-2022. This included 300 mutations associated with lineages including Alpha (28), B.1.351 (19), Gamma (32), Delta (25), Lambda (25), Mu (26), BA.1 (42), BA.2 (62), and BA.3 (41). Of these, 253 were unique mutations with 40,56,18 associated with BA.1, BA.2 and Delta respectively.

#### V-pipe

Reads were processed using V-pipe using the sars-cov-2 virus base config, which sets the aligner to bwa and the reference genome to the NCBI reference sequence NC_045512.2. V-pipe performed raw read quality filtering and alignment, and computed per-sample statistics, such as the per-position base counts. Detection of genomic variants in the mixed samples was performed using COJAC. For generating variant definitions, two different sources were used. A set of exhaustive lists of mutations by querying Cov-Spectrum for the Omicron variants BA.1 and BA.2, and Delta (B.1.617.2). For the Delta variant, mutations were imported from the mutation signature reported by Public Health England Genomics under the name “empathy-serve”. For COJAC and LolliPop, this “empathy-serve”-derived definition of Delta was used in addition to the exhaustive Omicron definitions. For COJAC, selected relevant amplicons were used covering combinations of mutations exclusive to B.1.617.2*, BA.1*, and BA.2*, by manual inspection, and then confirmed by querying Cov-Spectrum for the frequency at which these mutation combinations are found in the background. COJAC was run on the alignments produced by V-pipe on amplicons carrying cooccurrences of two or more mutations from the lists. For the detected variants, we quantified their relative abundances using LolliPop. LolliPop was run using the soft l1 loss, with default l1/l2 loss breakpoint at 0.1. LolliPop is tailored to analyzing time series wastewater sequencing data, and thus incorporates joint temporal kernel smoothing and deconvolution. However, in this benchmarking study the data was not a time series and hence smoothing was avoided by using a Gaussian kernel with bandwidth 10^-17^. ShoRAH was used to identify unknown lineages. Reads were merged paired-end using flash (version 1.2.11), before realigning to the reference using bwa-mem. The alignment was then tiled into the 98 ARTIC amplicon regions. For each amplicon region, local haplotypes were reconstructed by sampling from a Dirichlet Process Mixture Model. A Gibbs sampler was used to derive estimates of haplotypes and their frequencies in each amplicon region. The sampler was then run for a minimum of 300,000 steps or 15 times the read coverage. Only the last 100 sampling iterations for the final estimates were kept. To present the results, SNV calls were searched for each sample for mutations not found in any of the definitions (using the exhaustive mutation lists). The samples were sorted by descending number of such new mutations.

#### Freyja

Freyja preprocessing was performed by aligning paired-end FASTQ data to the reference genome with minimap2, followed by sorting and indexing of the resulting bam file with samtools. Bam files were then processed with the standard `freyja variants` and `freyja demix` commands (Freyja v1.3.7, doi:10.5281/zenodo.6585068), using the lineage barcodes from June 2, 2022.

#### Gromstole

FASTQ data were trimmed using cutadapt discarding trimmed reads below a minimum length of 10 (default) nucleotides. Then Minimap2 maps reads to the SARS-CoV-2 reference genome (Genbank accession NC_045512) using Gromstole with default settings. Constellations were v.0.1.10 with modification, which is forked cov-lineage constellations (May 5, 2022) with shared mutations between BA.1, BA.2.75, B.1.617.2, BA.4 and BA.5 removed, and BE.1 and BQ.1.1 added.

#### LCS

FASTQ files were loaded into LCS and processed with default settings and pre-generated markers (v1.2.124).

#### Lineagespot

Cleaned reads were mapped to the SARS-CoV-2 reference genome (Wuhan variant, NC_045512), using the Minimap2 tool. From this process, only the paired-end sequences were retained, while any other (unmatched, multiple mappings, etc.) were removed. In the next step, two different computational workflows were employed corresponding to the two different sequence lengths created by the NGS platforms. The first approach required primer removal, in which any primer sequences are excluded using the iVar tool. The final sequences are then remapped to the same reference genome. For the second approach, primer trimming and remapping to the reference genome is not applied. Finally, in order to be able to detect low frequency mutations, the freebayes variant caller was used with a low mutation frequency parameter of 0.01. Ultimately, all identified mutations were annotated using the SnpEf tool and the NC_045512.2 (version 5.0) database. The proposed variant calling pipeline (with all set parameters) is provided through the GitHub repository (https://github.com/BiodataAnalysisGroup/lineagespot/blob/master/inst/scripts/raw-data-analysis.md).

The annotated VCF files, corresponding to each sample from preprocessing, were given as input to lineagespot, together with the definition files of the target SARS-CoV-2 lineages’ profiles of lineages included in study (i.e. BA1, BA.2, Delta), as these were retrieved from outbreak.info. lineagespot ran for all samples altogether and the resulting tables were merged into a single file. Finally, the assignment of each sample to one of the provided options was performed by a semi-supervised assessment of each entry, in particular across samples that exhibited similar profiles (and metrics) across multiple definitions.

#### VLQ

Reference set construction was performed without temporal or geographical parameters due to the nature of the benchmark dataset, instead the reference set was created to be representative of all SARS-CoV-2 genomes in GISAID since the beginning of the pandemic until the time of download (June 12, 2022). In total 2527 sequences were selected using VLQ preprocessing (step 1), and the 34 reference genomes of the benchmark. Following reference set construction, the VLQ pipeline (steps 2 and 3) was run using default parameter settings.

#### Pipes *et al*

The reference database was built using the sarscov2 imputation command using 11,238 internal nodes from the GISAID SARS-CoV-2 global phylogeny of 7,603,548 genomes posted on May 1, 2022. Sequences from the internal nodes were estimated based on the maximum of the posterior probability of each nucleotide. The mismatch matrix was constructed using the eliminate strains program followed by EM_C_LLR.R (R-script) to run the EM-algorithm on the mismatch matrix to obtain the maximum likelihood estimates of the proportions of different haplotypes in a sample.

For tools/approaches that output relative abundances that do not sum to one, we classified the remaining relative abundance as ‘Other’ as they represent unclassified reads (Basic, Gromstole, alcov, LCS). For tools that predicted sublineages (alcov, basic, freyja, LCS and VLQ) of Delta, BA.1, or BA.2 we merged lineage abundances into their parent lineage, with remaining lineages classified as ‘Other’. Pipes et al. performed the merging of haplotypes and submitted the main-lineage distinction. The exception was for predictions used in evaluating the single lineage assignment for the synthetic and recombinant genomes (**Figure 3**) where the submitted lineages were kept for the tools: alcov, basic, freyja, LCS, and VLQ. In this instance we merged lineages only that belonged to one of the lineages in the Outbreak.info curated lineage list (2023-05-05).

## Acknowledgements

This study was supported by the Canadian Institutes for Health Research (CIHR) operating grant to the Coronavirus Variants Rapid Response Network (CoVaRR-Net). Data analyses were enabled by computing and storage resources provided by Compute Canada and Calcul Québec.

We gratefully acknowledge all data contributors, i.e., the Authors and their Originating laboratories responsible for obtaining the specimens, and their Submitting laboratories for generating the genetic sequence and metadata and sharing via the GISAID Initiative, on which this research is based.

## Supplementary Materials

**Supplementary Table 1.** The composition of each of the 100 benchmark samples. ‘lowcov’ indicates <50x depth and ‘hicov’ indicates 600x.

| Name | Lineages | Read length (bp) | Lineage ratio | Coverage |
| --- | --- | --- | --- | --- |
| Two lineages | BA.1:Delta | 250 | 75:25 | lowcov |
| Two lineages+Synth | BA.1:Delta:Synth | 250 | 65:25:10 | hicov |
| Two lineages | BA.1:Delta | 250 | 75:25 | hicov |
| Three lineages | BA.1:BA.2:Delta | 250 | 33:33:33 | lowcov |
| Two lineages+Synth | BA.1:Delta:Synth | 250 | 80:10:10 | lowcov |
| Two lineages | BA.1:Delta | 150 | 50:50 | lowcov |
| Three lineages+Synth | BA.1:BA.2:Delta:Synth | 250 | 30:30:30:10 | lowcov |
| Two lineages | BA.1:Delta | 250 | 1:99 | lowcov |
| Two lineages | BA.1:Delta | 250 | 90:10 | lowcov |
| Two lineages | BA.1:Delta | 250 | 90:10 | hicov |
| Three lineages+Synth | BA.1:BA.2:Delta:Synth | 250 | 10:10:70:10 | hicov |
| Two lineages (drop) | BA.1_drop:Delta | 250 | 25:75 | hicov |
| Two lineages | BA.1:Delta | 150 | 10:90 | lowcov |
| Two lineages (drop) | BA.1_drop:Delta | 250 | 90:10 | lowcov |
| Three lineages | BA.1:BA.2:Delta | 250 | 10:80:10 | lowcov |
| Two lineages (drop) | BA.1_drop:Delta | 250 | 1:99 | hicov |
| Single lineage | BA.2 | 250 | 100 | lowcov |
| Two lineages | BA.1:Delta | 150 | 75:25 | hicov |
| Two lineages | BA.1:BA.2 | 250 | 50:50 | hicov |
| Single lineage | BA.1 | 150 | 100 | hicov |
| Two lineages+Synth | BA.1:Delta:Synth | 250 | 1:89:10 | hicov |
| Single lineage | BA.1_drop | 250 | 100 | hicov |
| Single lineage | BA.1_drop | 150 | 100 | lowcov |
| Two lineages (drop) | BA.1_drop:Delta | 250 | 50:50 | hicov |
| Two lineages+Synth | BA.1:Delta:Synth | 250 | 65:25:10 | lowcov |
| Two lineages (drop) | BA.1_drop:Delta | 250 | 50:50 | lowcov |
| Two lineages (drop) | BA.1_drop:Delta | 250 | 75:25 | hicov |
| Two lineages (drop) | BA.1_drop:Delta | 250 | 1:99 | lowcov |
| Two lineages (drop) | BA.1_drop:Delta | 250 | 10:90 | hicov |
| Two lineages+Synth | BA.1:Delta:Synth | 250 | 1:89:10 | lowcov |
| Three lineages +Synth | BA.1:BA.2:Delta:Synth | 250 | 70:10:10:10 | hicov |
| Two lineages | BA.1:Delta | 150 | 25:75 | hicov |
| Two lineages | BA.1:BA.2 | 250 | 75:25 | hicov |
| Two lineages | BA.1:Delta | 150 | 99:1 | lowcov |
| Two lineages | BA.1:Delta | 150 | 25:75 | lowcov |
| Two lineages (drop) | BA.1_drop:Delta | 250 | 10:90 | lowcov |
| Two lineages+Synth | BA.1:Delta:Synth | 250 | 45:45:10 | lowcov |
| Two lineages (drop) | BA.1_drop:Delta | 250 | 25:75 | lowcov |
| Two lineages | BA.1:Delta | 150 | 10:90 | hicov |
| Single lineage | BA.1_drop | 150 | 100 | hicov |
| Two lineages | BA.1:Delta | 250 | 25:75 | lowcov |
| Three lineages+Synth | BA.1:BA.2:Delta:Synth | 250 | 10:70:10:10 | lowcov |
| Single lineage | BA.2 | 150 | 100 | lowcov |
| Three lineages | BA.1:BA.2:Delta | 250 | 10:10:80 | lowcov |
| Three lineages+Synth | BA.1:BA.2:Delta:Synth | 250 | 70:10:10:10 | lowcov |
| Two lineages | BA.1:Delta | 250 | 99:1 | hicov |
| Two lineages | BA.1:Delta | 150 | 50:50 | hicov |
| Two lineages | BA.1:Delta | 250 | 50:50 | lowcov |
| Two lineages | BA.1:Delta | 150 | 1:99 | lowcov |
| Two lineages | BA.1:Delta | 150 | 75:25 | lowcov |
| Single lineage | Delta | 250 | 100 | hicov |
| Single lineage | Recomb | 250 | 100 | lowcov |
| Two lineages | BA.1:Delta | 150 | 90:10 | lowcov |
| Single lineage | Delta | 150 | 100 | lowcov |
| Two lineages | BA.1:Delta | 250 | 10:90 | hicov |
| Two lineages | BA.1:Delta | 150 | 99:1 | hicov |
| Single lineage | Delta | 250 | 100 | lowcov |
| Two lineages | BA.1:Delta | 250 | 1:99 | hicov |
| Two lineages | BA.1:BA.2 | 250 | 50:50 | lowcov |
| Two lineages | BA.1:BA.2 | 250 | 25:75 | hicov |
| Two lineages | BA.1:Delta | 250 | 10:90 | lowcov |
| Two lineages+Synth | BA.1:Delta:Synth | 250 | 25:65:10 | hicov |
| Single lineage | BA.1 | 150 | 100 | lowcov |
| Two lineages | BA.1:Delta | 150 | 1:99 | hicov |
| Three lineages | BA.1:BA.2:Delta | 250 | 80:10:10 | lowcov |
| Single lineage | Synth | 250 | 100 | lowcov |
| Three lineages | BA.1:BA.2:Delta | 250 | 33:33:33 | hicov |
| Single lineage | Synth | 150 | 100 | hicov |
| Two lineages (drop) | BA.1_drop:Delta | 250 | 75:25 | lowcov |
| Two lineages | BA.1:Delta | 150 | 90:10 | hicov |
| Single lineage | BA.1_drop | 250 | 100 | lowcov |
| Single lineage | Delta | 150 | 100 | hicov |
| Two lineages+Synth | BA.1:Delta:Synth | 250 | 10:80:10 | lowcov |
| Two lineages | BA.1:Delta | 250 | 99:1 | lowcov |
| Single lineage | Synth | 250 | 100 | hicov |
| Single lineage | BA.1 | 250 | 100 | hicov |
| Two lineages (drop) | BA.1_drop:Delta | 250 | 99:1 | lowcov |
| Two lineages+Synth | BA.1:Delta:Synth | 250 | 10:80:10 | hicov |
| Three lineages+Synth | BA.1:BA.2:Delta:Synth | 250 | 10:70:10:10 | hicov |
| Two lineages | BA.1:BA.2 | 250 | 75:25 | lowcov |
| Two lineages | BA.1:Delta | 250 | 25:75 | hicov |
| Three lineages | BA.1:BA.2:Delta | 250 | 80:10:10 | hicov |
| Single lineage | Recomb | 250 | 100 | hicov |
| Two lineages | BA.1:BA.2 | 250 | 25:75 | lowcov |
| Two lineages+Synth | BA.1:Delta:Synth | 250 | 45:45:10 | hicov |
| Two lineages (drop) | BA.1_drop:Delta | 250 | 99:1 | hicov |
| Two lineages+Synth | BA.1:Delta:Synth | 250 | 80:10:10 | hicov |
| Two lineages | BA.1:Delta | 250 | 50:50 | hicov |
| Two lineages+Synth | BA.1:Delta:Synth | 250 | 89:1:10 | hicov |
| Single lineage | BA.2 | 150 | 100 | hicov |
| Two lineages+Synth | BA.1:Delta:Synth | 250 | 25:65:10 | lowcov |
| Three lineages | BA.1:BA.2:Delta | 250 | 10:80:10 | hicov |
| Three lineages+Synth | BA.1:BA.2:Delta:Synth | 250 | 10:10:70:10 | lowcov |
| Three lineages+Synth | BA.1:BA.2:Delta:Synth | 250 | 30:30:30:10 | hicov |
| Two lineages+Synth | BA.1:Delta:Synth | 250 | 89:1:10 | lowcov |
| Two lineages (drop) | BA.1_drop:Delta | 250 | 90:10 | hicov |
| Single lineage | BA.1 | 250 | 100 | lowcov |
| Single lineage | Synth | 150 | 100 | lowcov |
| Three lineages | BA.1:BA.2:Delta | 250 | 10:10:80 | hicov |
| Single lineage | BA.2 | 250 | 100 | hicov |

**Supplementary Table 2.**
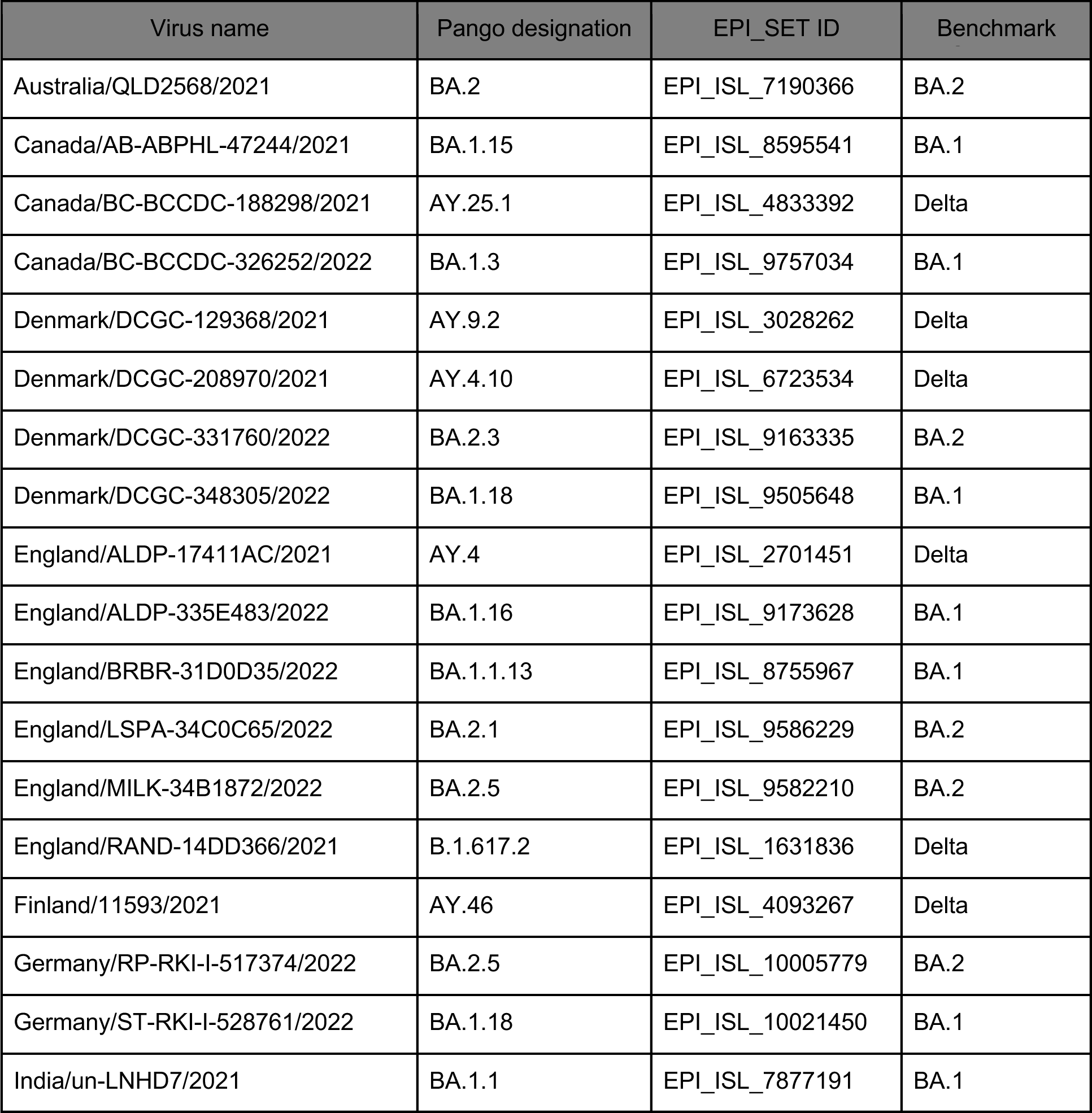
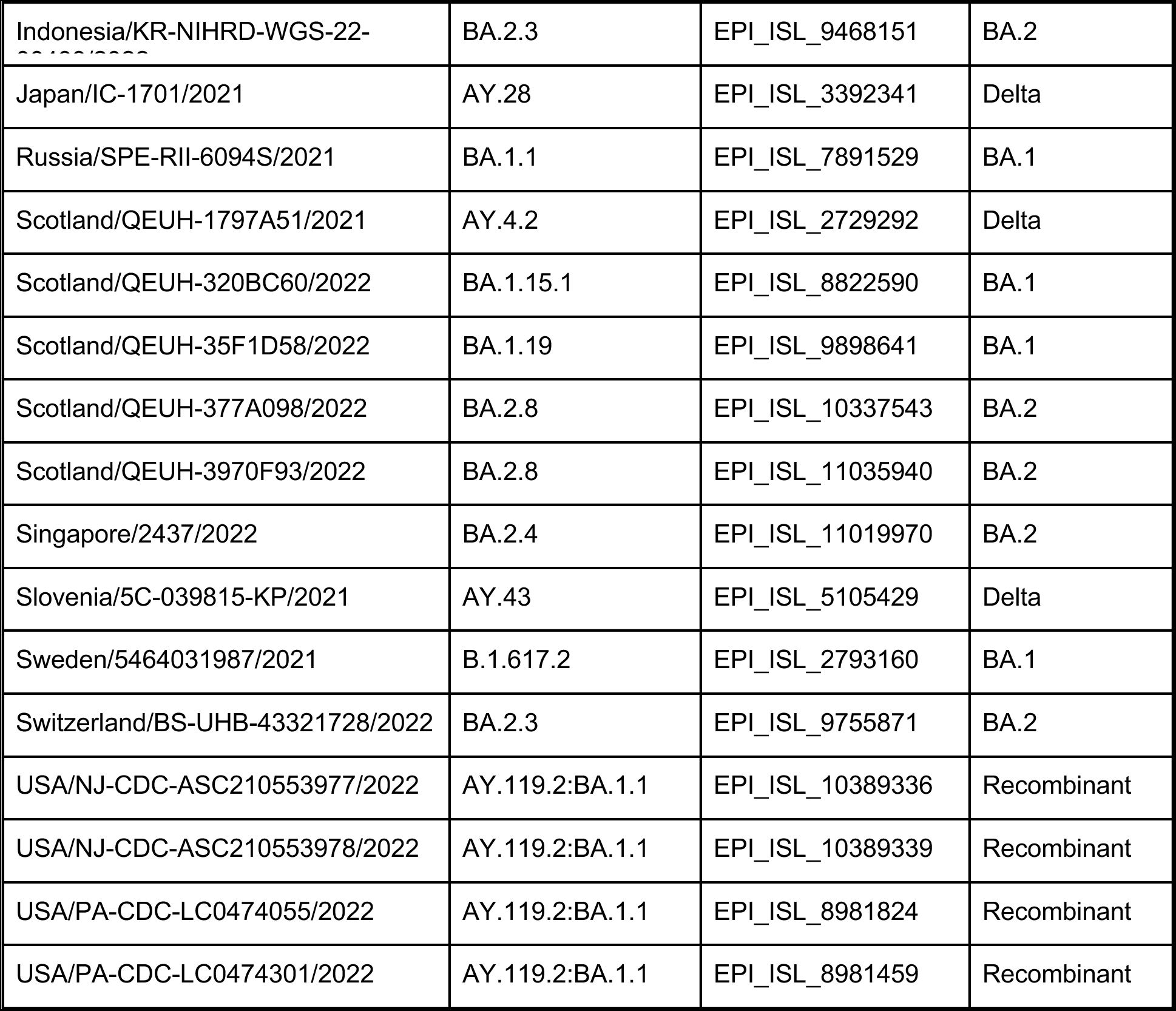
Genomes from GISAID used in building the benchmark dataset available on 2022-03-06 via gisaid.org/EPI_SET_230802sn.

## GISAID Data Availability

GISAID Identifier: EPI_SET_230802sn doi: 10.55876/gis8.230802sn

All genome sequences and associated metadata in this dataset are published in GISAID’s EpiCoV database. To view the contributors of each individual sequence with details such as accession number, Virus name, Collection date, Originating Lab and Submitting Lab and the list of Authors, visit 10.55876/gis8.230802sn

## GISAID Data Snapshot

- EPI_SET_230802sn is composed of 34 individual genome sequences.
- The collection dates range from 2021-04-06 to 2022-03-06;
- Data were collected in 15 countries and territories;
- All sequences in this dataset are compared relative to hCoV-19/Wuhan/WIV04/2019 (WIV04), the official reference sequence employed by GISAID (EPI_ISL_402124). Learn more at https://gisaid.org/WIV04.

## References

1. Choi, P. M. et al. Wastewater-based epidemiology biomarkers: Past, present and future. TrAC Trends Anal. Chem. 105, 453–469 (2018).

2. Sims, N. & Kasprzyk-Hordern, B. Future perspectives of wastewater-based epidemiology: Monitoring infectious disease spread and resistance to the community level. Environ. Int. 139, 105689 (2020).

3. Naughton, C. C., et al. Show us the data: global COVID-19 wastewater monitoring efforts, equity, and gaps. FEMS Microbes 4, xtad003 (2023).

4. Tang, A. et al. Detection of Novel Coronavirus by RT-PCR in Stool Specimen from Asymptomatic Child, China. Emerg. Infect. Dis. 26, 1337–1339 (2020).

5. Chen, Y. et al. The presence of SARS-CoV-2 RNA in the feces of COVID-19 patients. J. Med. Virol. 92, 833–840 (2020).

6. Wu, Y. et al. Prolonged presence of SARS-CoV-2 viral RNA in faecal samples. Lancet Gastroenterol. Hepatol. 5, 434–435 (2020).

7. Xiao, F. et al. Infectious SARS-CoV-2 in feces of patient with severe COVID-19. Emerg. Infect. Dis. 26, 1920–1922 (2020).

8. Ciannella, S., González-Fernández, C. & Gomez-Pastora, J. Recent progress on wastewater-based epidemiology for COVID-19 surveillance: A systematic review of analytical procedures and epidemiological modeling. Sci. Total Environ. 878, 162953 (2023).

9. Wolfe, M. et al. Detection of SARS-CoV-2 variants Mu, Beta, Gamma, Lambda, Delta, Alpha, and Omicron in wastewater settled solids using mutation-specific assays is associated with regional detection of variants in clinical samples. Appl. Environ. Microbiol. 88, e00045–22 (2022).

10. Karthikeyan, S. et al. Wastewater sequencing reveals early cryptic SARS-CoV-2 variant transmission. Nature 609(7925), 101–108 (2022).

11. Xu, X. et al. Real-time allelic assays of SARS-CoV-2 variants to enhance sewage surveillance. Water Res. 220, 118686 (2022).

12. Cancela, F. et al. Wastewater surveillance of SARS-CoV-2 genomic populations on a country-wide scale through targeted sequencing. PLOS ONE 18, e0284483 (2023).

13. Kuroiwa, M. et al. Targeted amplicon sequencing of wastewater samples for detecting SARS-CoV-2 variants with high sensitivity and resolution. Sci. Total Environ. 893, 164766 (2023).

14. La Rosa, G. et al. Wastewater surveillance of SARS-CoV-2 variants in October– November 2022 in Italy: detection of XBB.1, BA.2.75 and rapid spread of the BQ.1 lineage. Sci. Total Environ. 873, 162339 (2023).

15. Sutton, M. et al. Detection of SARS-CoV-2 B.1.351 (Beta) variant through wastewater surveillance before case detection in a community, Oregon, USA. Emerg. Infect. Dis. 28, (2022).

16. Manuel, D., Amadei, C. A., Campbell, J. R., Brault, J.-M. & Veillard, J. Strengthening public health surveillance through wastewater testing. (2022).

17. Wölfel, R. et al. Virological assessment of hospitalized patients with COVID-2019. Nature 581, 465–469 (2020).

18. Trottier, J. et al. Post-lockdown detection of SARS-CoV-2 RNA in the wastewater of Montpellier, France. One Health 10, 100157 (2020).

19. McMinn, B. R. et al. Assessment of two volumetrically different concentration approaches to improve sensitivities for SARS-CoV-2 detection during wastewater monitoring. J. Virol. Methods 311, 114645 (2023).

20. Wigginton, K. R., Ye, Y. & Ellenberg, R. M. Emerging investigators series: the source and fate of pandemic viruses in the urban water cycle. Environ. Sci. Water Res. Technol. 1, 735–746 (2015).

21. Parra-Arroyo, L. et al. Degradation of viral RNA in wastewater complex matrix models and other standards for wastewater-based epidemiology: A review. TrAC Trends Anal. Chem. 158, 116890 (2023).

22. Bertels, X. et al. Factors influencing SARS-CoV-2 RNA concentrations in wastewater up to the sampling stage: A systematic review. Sci. Total Environ. 820, 153290 (2022).

23. Wu, F. et al. Making waves: Wastewater surveillance of SARS-CoV-2 in an endemic future. Water Res. 219, 118535 (2022).

24. Agrawal, S., Orschler, L. & Lackner, S. Metatranscriptomic analysis reveals SARS-CoV-2 mutations in wastewater of the Frankfurt Metropolitan Area in Southern Germany. Microbiol. Resour. Announc. 10, (2021).

25. Novoa, B. et al. Wastewater and marine bioindicators surveillance to anticipate COVID-19 prevalence and to explore SARS-CoV-2 diversity by next generation sequencing: One-year study. Sci. Total Environ. 833, 155140 (2022).

26. Smith, M. F. et al. Baseline sequencing surveillance of public clinical testing, hospitals, and community wastewater reveals rapid emergence of SARS-CoV-2 Omicron variant of concern in Arizona, USA. mBio 14, e03101–22 (2023).

27. Rios, G. et al. Monitoring SARS-CoV-2 variants alterations in Nice neighborhoods by wastewater nanopore sequencing. Lancet Reg. Health - Eur. 10, 100202 (2021).

28. Ellmen, I., et al. Alcov: Estimating Variant of Concern Abundance from SARS-CoV-2 Wastewater Sequencing Data. medRxiv 2021.06.03.21258306 (2021).

29. Chen, C. et al. CoV-Spectrum: analysis of globally shared SARS-CoV-2 data to identify and characterize new variants. Bioinformatics 38, 1735–1737 (2022).

30. N’Guessan, A. et al. Detection of prevalent SARS-CoV-2 variant lineages in wastewater and clinical sequences from cities in Québec, Canada. medRxiv 2022.02.01.22270170 (2022).

31. Poon, A., Becker, Devin & Gugan, Gopi. Gromstole. Github repository https://github.com/PoonLab/gromstole (2023).

32. Valieris, R. et al. A mixture model for determining SARS-Cov-2 variant composition in pooled samples. Bioinformatics 38, 1809–1815 (2022).

33. Pechlivanis, N. et al. Detecting SARS-CoV-2 lineages and mutational load in municipal wastewater and a use-case in the metropolitan area of Thessaloniki, Greece. Sci. Rep. 12, 2659 (2022).

34. Pipes, L., Chen, Z., Afanaseva, S. & Nielsen, R. Estimating the relative proportions of SARS-CoV-2 haplotypes from wastewater samples. *Cell Rep*. Methods 2, 100313 (2022).

35. Baaijens, J. A. et al. Lineage abundance estimation for SARS-CoV-2 in wastewater using transcriptome quantification techniques. Genome Biol. 23, 236 (2022).

36. Bray, N. L., Pimentel, H., Melsted, P. & Pachter, L. Near-optimal probabilistic RNA-seq quantification. Nat. Biotechnol. 34, 525–527 (2016).

37. Posada-Céspedes, S. et al. V-pipe: a computational pipeline for assessing viral genetic diversity from high-throughput data. Bioinformatics 37, 1673–1680 (2021).

38. Jahn, K. et al. Early detection and surveillance of SARS-CoV-2 genomic variants in wastewater using COJAC. Nat. Microbiol. 7(8) 1151–1160 (2022).

39. David Dreifuss, Ivan Topolsky, Pelin Icer Baykal, & Niko Beerenwinkel. Tracking SARS-CoV-2 genomic variants in wastewater sequencing data with LolliPop. medRxiv 2022.11.02.22281825 (2022).

40. Zagordi, O., Bhattacharya, A., Eriksson, N. & Beerenwinkel, N. ShoRAH: estimating the genetic diversity of a mixed sample from next-generation sequencing data. BMC Bioinformatics 12, 119 (2011).

41. O’Reilly, K. M., Allen, D. J., Fine, P. & Asghar, H. The challenges of informative wastewater sampling for SARS-CoV-2 must be met: lessons from polio eradication. Lancet Microbe 1, e189–e190 (2020).

42. Shaw, A. G. et al. Rapid and sensitive direct detection and identification of poliovirus from stool and environmental surveillance samples by use of Nanopore sequencing. J. Clin. Microbiol. 58, (2020).

43. Ahmed, W. et al. First confirmed detection of SARS-CoV-2 in untreated wastewater in Australia: A proof of concept for the wastewater surveillance of COVID-19 in the community. Sci. Total Environ. 728, 138764 (2020).

44. Quick, J. nCoV-2019 sequencing protocol v3 (LoCost). (2020) doi:10.17504/protocols.io.bp2l6n26rgqe/v3.

45. 45. Liu, T., et al. A benchmarking study of SARS-CoV-2 whole-genome sequencing protocols using COVID-19 patient samples. iScience 24, (2021).

46. Public Health Agency of Canada. Wastewater sequencing trend report: Detection of SARS-CoV-2 variants of concern by metagenomic sequencing. https://www.canada.ca/en/public-health/services/emergency-preparedness-response/wastewater-monitoring.html (2023).

47. Hart, O. E. & Halden, R. U. Computational analysis of SARS-CoV-2/COVID-19 surveillance by wastewater-based epidemiology locally and globally: Feasibility, economy, opportunities and challenges. Sci. Total Environ. 730, 138875 (2020).

48. Dharmadhikari, T. et al. High throughput sequencing based direct detection of SARS-CoV-2 fragments in wastewater of Pune, West India. Sci. Total Environ. 807, 151038 (2022).

49. Libuit, K., et al. Quality control solutions for SARS-CoV-2 genomic analysis. Public Health Alliance for Genomic Epidemiology https://pha4ge.org/resource/qc-solutions-for-sars-cov-2-genomic-analysis/ (2022).

50. Boehm, A. B. et al. Wastewater surveillance of human influenza, metapneumovirus, parainfluenza, respiratory syncytial virus (RSV), rhinovirus, and seasonal coronaviruses during the COVID-19 pandemic. medRxiv. 2022.09.22.22280218 (2023).

51. Tisza, M. et al. Wastewater sequencing reveals community and variant dynamics of the collective human virome. Nat. Commun. 14, 6878 (2023).

52. Li, J. et al. A global aircraft-based wastewater genomic surveillance network for early warning of future pandemics. Lancet Glob. Health 11, e791–e795 (2023).

53. Medina, C. Y. et al. The need of an environmental justice approach for wastewater based epidemiology for rural and disadvantaged communities: A review in California. Curr. Opin. Environ. Sci. Health 27, 100348 (2022).

54. Khare, S., et al. GISAID’s role in pandemic response. China CDC Wkly. 3, 1049–1051 (2021).

55. Lacek, K. A. et al. SARS-CoV-2 Delta–Omicron Recombinant Viruses, United States. Emerg. Infect. Dis. 28, (2022).

56. Shen, W., Le, S., Li, Y. & Hu, F. SeqKit: A Cross-Platform and ultrafast toolkit for FASTA/Q file manipulation. PLOS ONE 11, e0163962 (2016).

57. Gourlé, H., Karlsson-Lindsjö, O., Hayer, J. & Bongcam-Rudloff, E. Simulating Illumina metagenomic data with InSilicoSeq. Bioinformatics 35, 521–522 (2019).

